# Proximity labeling reveals dynamic changes in the SQSTM1 protein network

**DOI:** 10.1101/2023.12.12.571324

**Authors:** Alejandro N. Rondón Ortiz, Lushuang Zhang, Peter E.A. Ash, Avik Basu, Sambhavi Puri, Sophie J.F. van der Spek, Zihan Wang, Luke Dorrian, Andrew Emili, Benjamin Wolozin

## Abstract

Sequestosome1 (SQSTM1) is an autophagy receptor that mediates degradation of intracellular cargo, including protein aggregates, through multiple protein interactions. These interactions form the SQSTM1 protein network, and these interactions are mediated by SQSTM1 functional interaction domains, which include LIR, PB1, UBA and KIR. Technological advances in cell biology continue to expand our knowledge of the SQSTM1 protein network and of the relationship of the actions of the SQSTM1 protein network in cellular physiology and disease states. Here we apply proximity profile labeling to investigate the SQSTM1 protein interaction network by fusing TurboID with the human protein SQSTM1 (TurboID::SQSTM1). This chimeric protein displayed well-established SQSTM1 features including production of SQSTM1 intracellular bodies, binding to known SQSTM1 interacting partners, and capture of novel SQSTM1 protein interactors. Strikingly, aggregated tau protein altered the protein interaction network of SQSTM1 to include many stress-associated proteins. We demonstrate the importance of the PB1 and/or UBA domains for binding network members, including the K18 domain of tau. Overall, our work reveals the dynamic landscape of the SQSTM1 protein network and offers a resource to study SQSTM1 function in cellular physiology and disease state.

## INTRODUCTION

Sequestosome 1 (SQSTM1), commonly referred to as p62, is a selective autophagy receptor that recognizes and targets specific cellular cargo for degradation, such as misfolded proteins and damaged organelles (1–3). SQSTM1 acts as an adaptor that couples cargo to the Microtubule Associated Protein 1 Light Chain 3β (LC3B) protein (and other proteins from the ATG8, family LC3A, LC3C, GABARAP, GABARAPL1 and GABARAPL2) for transport through the autophagic system for degradation(3–6). In addition to its role in autophagy, SQSTM1 participates in other cellular processes including signal transduction and cellular stress response (7–10). These cellular processes involve dynamic interactions via multiple discrete SQSTM1 functional domains: Phox and Bem1 (PB1), ZZ-type zinc finger, TRAF-6 binding domain, KEAP1-binding region (KIR), LC3-interacting region (LIR) and the ubiquitin-associated domain (UBA). SQSTM1 protein intracellular localization is predominantly cytosolic, but SQSTM1 has also been detected in the nuclear compartment and its nuclear localization has been implicated as a nucleus-to-cytosol shuttle for ubiquitinylated nuclear proteins(11).

Proteins often operate as part of complex molecular networks orchestrating a plethora of biological processes (12, 13). The SQSTM1 protein network exhibits central roles in both cellular proteostasis and pathology (14–16). The importance of SQSTM1 in catabolic processes (*i.e.* autophagy and proteostasis) is pivotal for the understanding of disorders such as neurodegenerative disorders, cancers, and metabolic syndromes (17–20). Thus, characterization of the dynamic SQSTM1 protein interaction network can be used as a tool to probe disease processes broadly, identifying cargo proteins that are ubiquitinated, misfolded or aggregated. Elucidating SQSTM1 protein networks in normal and disease states could provide insights into the evolution of pathological processes, broadening the comprehension of cellular mechanisms underlying proteostasis, and potentially identifying novel disease markers and therapeutic targets.

Targeted proximity labelling has emerged as an innovative proteomic strategy to explore protein-protein interactions. Unbiased enzyme-based approaches confer the advantage of capturing macromolecular interactions within a near native spatial and temporal cellular milieu (13). Notably, fusion of the engineered biotin ligase TurboID to a target protein confers biotinylation of other physically proximal proteins which can subsequently be identified by tandem mass spectrometry (21–24).

In this study, we generated and characterized TurboID-based SQSTM1 chimeric proteins to probe SQSTM1 dynamics in native and pathological contexts. We identify multiple novel SQSTM1 network members and demonstrate that the SQSTM1 PB1 domain is necessary for interaction of SQSTM1 with the aggregation prone K18-Tau peptide. The resulting networks reveal large-scale remodeling of SQSTM1 protein cargo in response to pathological states.

## RESULTS

### Proximity labeling of SQSTM1 protein network

The autophagy receptor SQSTM1 interacts with many proteins under both basal and stress conditions. These interactions take place through the SQSTM1 interacting functional domains that include LIR, PB1, UBA and KIR (1, 25, 26). Moreover, novel SQSTM1 interacting partners continue to be discovered that aim to describe SQSTM1-associated biological process (27–29). To enhance the understanding of the SQSTM1 protein network, we applied a systematic approach centered on proximity labeling and generated a chimeric FLAG-tagged human full-length TurboID::SQSTM1 (22, 23, 30).

To test functionality, HEK-293T cells were transduced with lentiviruses (LVs, multiplicity of infection = 2) expressing either FLAG::TurboID (non-specific background control) or FLAG::TurboID::SQSTM1 (Figure 1A). Consistent with expectation, immunofluorescence analysis of the FLAG epitope demonstrated that FLAG::TurboID::SQSTM1 fusion formed intracellular SQSTM1 bodies, a characteristic feature of the native SQSTM1 protein (Figure 1B). In contrast, FLAG::TurboID control protein displayed a dispersed intracellular expression pattern, similar to that observed in prior studies (3, 31). Imaging the enzymatic biotinylating activity of the FLAG::TurboID::SQSTM1 fusion construct showed a pattern of biotinylation that selectively co-localized with SQSTM1 positive structures (Figure 1B). In contrast, the FLAG::TurboID control protein exhibited a dispersed pattern, similar to that observed with the anti-FLAG antibody (Figure 1B).

**Figure 1.**
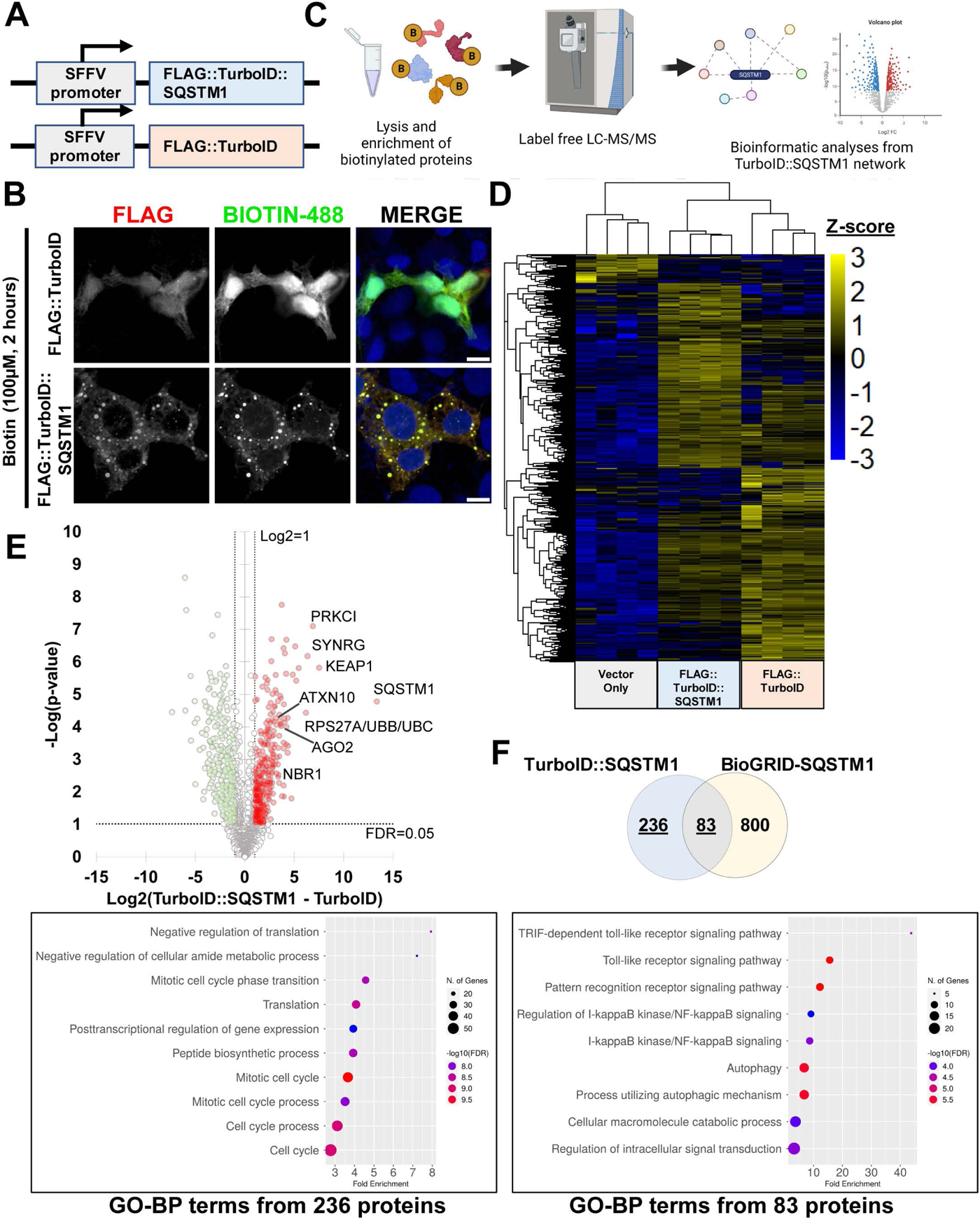
Proximity labeling of SQSTM1 networks. A) Outline of the lentiviral constructs FLAG::TurboID and FLAG::TurboID::SQSTM1. B) Immunofluorescence images of HEK-293T cells transduced with either FLAG::TurboID or FLAG::TurboID::SQSTM1 constructs (scale bar = 10µm). C) Outline of the targeted proximity labeling proteomic approach. D) Hierarchical clustering by Z-score intensities of differentially selected proteins in transduced HEK-293T cells. Proteins were selected using ANOVA and permutation-based FDR <0.05 (n=4). Data listed in Supplementary Table 1, Sheet-1. E) Quantitative comparison of HEK-293T cells transduced-with either FLAG::TurboID or FLAG::TurboID::SQSTM1 constructs using a Log2 FC(TurboID::SQSTM1 – TurboID) of 1 and FDR < 0.05. (n=4). Known and novel SQSTM1 protein interacting partners are highlighted in the volcano-plot. Data listed in Supplementary Table 1, Sheet-2. F) Comparison of significantly enriched proteins from SQSTM1-TurboID dataset (Figure 1E) with BioGRID-SQSTM1 dataset. GO-BP, Gene Ontology – Biological processes; FDR, False Discovery Rate; FC, Fold-Change.

We used Streptavidin-loaded magnetic beads to pull down labelled proteins from LVs-transduced HEK-293T cells expressing either vector-only (Vo), FLAG::TurboID::SQSTM1 or FLAG::TurboID constructs. We subsequently identified labeled proteins by high performance liquid-chromatography coupled to Orbitrap tandem mass spectrometry (LC-MS/MS, Figure 1C). Following the precision proteomics analysis, we used hierarchical clustering to reveal proteins that were labeled preferentially by FLAG::TurboID::SQSTM1 compared to FLAG::TurboID (Figure 1D and Supporting Information-1, Sheet-1). Quantitative comparisons demonstrated that FLAG::TurboID::SQSTM1 enriched a diverse array of 319 cellular proteins, including known SQSTM1 interacting partners like KEAP1, PRKCI, NBR1, Ubiquitin and others (Figure 1E and Supporting Information-1, Sheet-2).

We cross-referenced with previously annotated interactors comprising the BioGRID-SQSTM1 dataset (32). This overlap highlighted 83 proteins between these two interactome datasets. In addition, FLAG::TurboID::SQSTM1 construct labeled an additional 236 novel SQSTM1-interacting proteins that were not previously annotated (Figure 1F, Venn diagram). Functional enrichment analysis of gene ontology annotation terms associated with the 236 unique proteins demonstrated an enrichment for factors linked to translation and the cell cycle (Figure 1F, left panel), whereas the overlapping subset of 83 proteins were associated with autophagy and catabolic processes typically associated with core SQSTM1 biological functions (Figure 1F, right panel).

### SQSTM1 proximity labeling identifies known and novel SQSTM1 interactors

To validate some of the interactors from the SQSTM1 network recovered by FLAG::TurboID::SQSTM1 (Figures 1D and 1E), we performed independent co-immunoprecipitation experiments. After transducing HEK-293T cells with LVs expressing either Vector-only, FLAG::TurboID or FLAG::TurboID::SQSTM1, biotinylated proteins were pulled-down and resolved by immunoblotting. As seen in Figure 2A, endogenous KEAP1, endogenous SQSTM1 and ubiquitin were detected preferentially with FLAG::TurboID::SQSTM1 compared to the FLAG::TurboID control.

**Figure 2.**
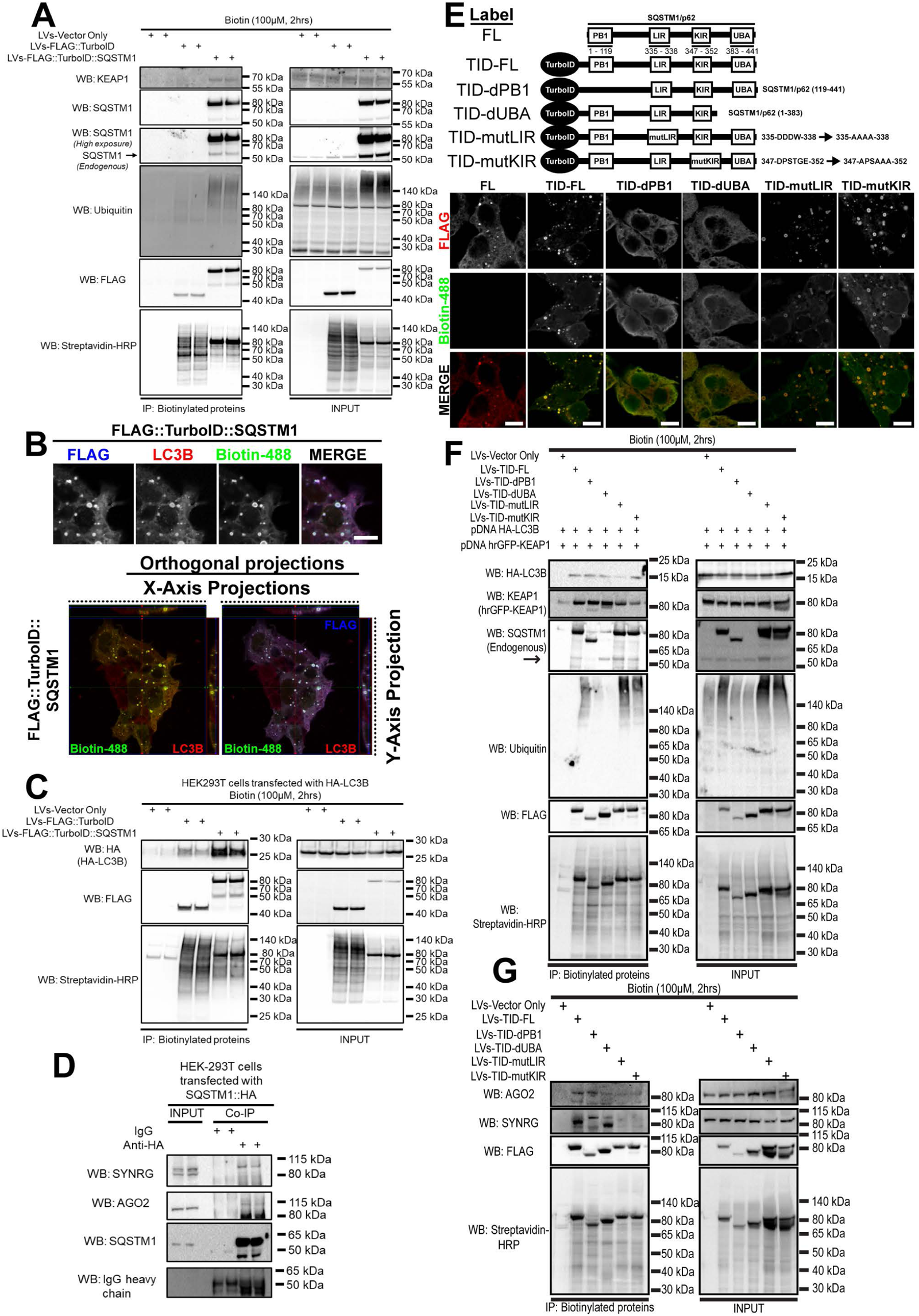
SQSTM1 proximity labelling identifies known and novel SQSTM1 interactors. A) Pull-down of biotinylated proteins from transduced HEK-293T cells. Eluted proteins were detected by Western blot, and it demonstrates the interaction of TurboID::SQSTM1 with KEAP1, endogenous SQSTM1 and Ubiquitin. B) Endogenous LC3B immunofluorescence images of HEK-293T cells transduced with TurboID::SQSTM1 construct. Lower panel contains orthogonal projections indicating that FLAG::TurboID::SQSTM1 expression, and biotinylated proteins, co-localized with endogenous LC3B (scale bar = 10µm). C) HA::LC3B interaction with TurboID::SQSTM1 detected by pull-down of biotinylated proteins. HEK-293T cells were previously transduced with TurboID and TurboID::SQSTM1 and then transfected with a HA::LC3B plasmid. D) SYNRG and AGO2 interaction with SQSTM1::HA as determined by HA tag co-immunoprecipitation using HEK-293T cells. E) Schematic representation of the SQSTM1 domain-function analysis from TurboID::SQSTM1 iterations. Immunofluorescence analysis of the different TurboID::SQSTM1 iterations demonstrate that deletion of both PB1 and UBA domains are essential for SQSTM1 bodies assembly. F) Pull-down of biotinylated proteins from HEK-293T cells transduced with the different iterations of TurboID::SQSTM1 constructs. These cells were transiently co-transfected with plasmids that encode for HA::LC3B and hrGFP::KEAP1. Pull-down of biotinylated proteins demonstrated the key roles of SQSTM1 domains to interact with known SQSTM1-interacting partners. G) Pull-down of biotinylated proteins from cells transduced with TurboID::SQSTM1 iterations demonstrated that AGO2-SQSTM1 interaction requires LIR, KIR and UBA domains. Whereas SYNRG-SQSTM1 interaction only occurs with the full-length SQSTM1 protein.

SQSTM1 links ubiquitinylated proteins to autophagosome-anchored proteins (*e.g.* LC3B, GABARAP and other members from the ATG8 gene family) through its LIR region (5, 33, 34). To test whether FLAG::TurboID::SQSTM1 preserves an active LIR domain, both endogenous and ectopically expressed LC3B protein were monitored in HEK-293T cells by immunofluorescence and streptavidin pull-down, respectively. Orthogonal projections demonstrated that both FLAG::TurboID::SQSTM1 and endogenous LC3B co-localized in the same compartment (Figure 2B). In addition, HEK-293T cells expressing either TurboID or TurboID::SQSTM1 were transfected with HA::LC3B, biotinylated proteins were pulled-down, and the HA-tag (HA::LC3B) was then determined by Western-blot. As expected, the HA::LC3B-TurboID::SQSTM1 interaction was greater compared to the HA::LC3B and TurboID interaction (Figure 2C). These observations suggest that FLAG::TurboID::SQSTM1 preserves its SQSTM1 functional domains PB1, LIR, KIR, and UBA.

The SQSTM1 interactome dataset (Figure 1F) also identified 236 previously unknown SQSTM1 interacting proteins. From this dataset, we selected the proteins, Synergin γ (SYNRG) and Argonaut 2 (AGO2) for validation based on their strong signals (Figure 1E). To corroborate these interactions, we immune-precipitated soluble protein lysates from HEK-293T cells transfected with a plasmid expressing epitope HA-tagged SQSTM1, and analyzed the co-purifying proteins by Western-blot. Both SYNRG and AGO2 co-purified with SQSTM1 (Figure 2D). Thus, our proximity proteomics screen identified previously unknown SQSTM1 protein interactors in HEK-293T cells.

To complement the FLAG::TurboID::SQSTM1 proximity labeling approach, we designed an array of TurboID::SQSTM1 constructs containing either truncated protein sequences or loss-of-function mutations to inhibit SQSTM1 functional protein domains (5, 35, 36). We truncated the PB1 domain (TID-dPB1, SQSTM1 deletion of 1-118) and the UBA domain (TID-dUBA, SQSTM1 deletion of 383 – 441). We also generated dysfunctional LIR region (TID-mutLIR, SQSTM1 335-DDDW-338 → 335-AAAA-338), and dysfunctional KIR region (TID-mutKIR, SQSTM1 347-DPSTGE-352 → 347-APSAAA-352), as observed in Figure 2E.

Deletion of either the PB1 or UBA domain in the TurboID::SQSTM1 construct abolished SQSTM1 bodies (Figure 2E), as previously reported (37). In addition, the protein-interaction profile of these TurboID::SQSTM1 constructs were monitored by pull-down of biotinylated proteins and Western blot. As predicted, truncated versions and loss-of-function mutations reduce the interaction of these constructs with known SQSTM1 interacting partners, HA::LC3B, GFP::Keap1, endogenous SQSTM1 and endogenous ubiquitinylated proteins (Figure 2F). Thus, these TurboID::SQSTM1 constructs can be exploited to study interaction partners that depend on these SQSTM1 functional domains. AGO2 and SYNRG were selected from our SQSTM1 interactome dataset (Figure 1F). Next, the interactions of SQSTM1 with these two proteins were validated using co-immunoprecipitation (Figure 2D). We evaluated the array of TurboID::SQSTM1 dysfunctional-domains constructs using HEK-293T cells to determine the SQSTM1 functional domains responsible for interaction with these two proteins. Subsequently, biotinylated protein pull-down was analyzed by immunoblotting. The results demonstrate that the AGO2-SQSTM1 interaction requires LIR, KIR and UBA domains, whereas the SYNRG-SQSTM1 interaction only occurs with the full-length SQSTM1 protein (Figure 2G). These results demonstrate the utility of these TurboID::SQSTM1 constructs to explore the SQSTM1 interaction network and determine the domain requirements for interaction of SQSTM1 with its binding partners.

### Effect of TurboID::SQSTM1 on autophagy flux

Our observations demonstrate that the TurboID::SQSTM1 protein interacts with essential proteins of the proteostasis machinery (Ubiquitin, LC3B, endogenous SQSTM1 protein, and others; as seen in Figures 1 and 2). SQSTM1 protein is known to serve as a bridge between degradation of intracellular cargo and the autophagy machinery. It was reported that SQSTM1 over-expression stalls autophagic flux (38), so we measured autophagic flux to evaluate the activity of TurboID::SQSTM1 chimeric protein on the autophagy pathway. HEK-293T cells were transduced with LVs expressing the autophagy flux sensor mCherry::EGFP::LC3B. This widely used sensor emits red and green fluorescence at neutral pH, but in acidic pH (lysosomal and autophagosome compartments) green fluorescence is dramatically reduced (39–41). Thus, the EGFP/mCherry ratio is an indicator of the autophagy flux.

HEK-293T cells expressing mCherry::EGFP::LC3B were transduced with a lentiviral-particles (MOI of 2) expressing either Vector-only, FLAG::TurboID::SQSTM1 or FLAG::SQSTM1 proteins. Next, these cells were starved using HBSS ± bafilomycin A1 (100nM, 4 hours); the bafilomycin was used as a positive control for blockade of autophagic flux. Following the treatment, the cells were fixed, stained for FLAG-tag and fluorescence intensity was monitored by confocal microscopy. As expected, overexpression of both FLAG::TurboID::SQSTM1 and FLAG::SQSTM1 inhibited autophagic flux (Figure 3A, 3B, and Supporting Information-1, Sheet-9), with the signal from mCherry::EGFP::LC3B accumulating coincident with the FLAG signal. Overall, these results show that overexpression of both FLAG::TurboID::SQSTM1 and FLAG::SQSTM1 inhibit the autophagy flux in HEK-293T cells, and that FLAG::TurboID::SQSTM1 functions in a manner similar to FLAG::SQSTM1.

**Figure 3.**
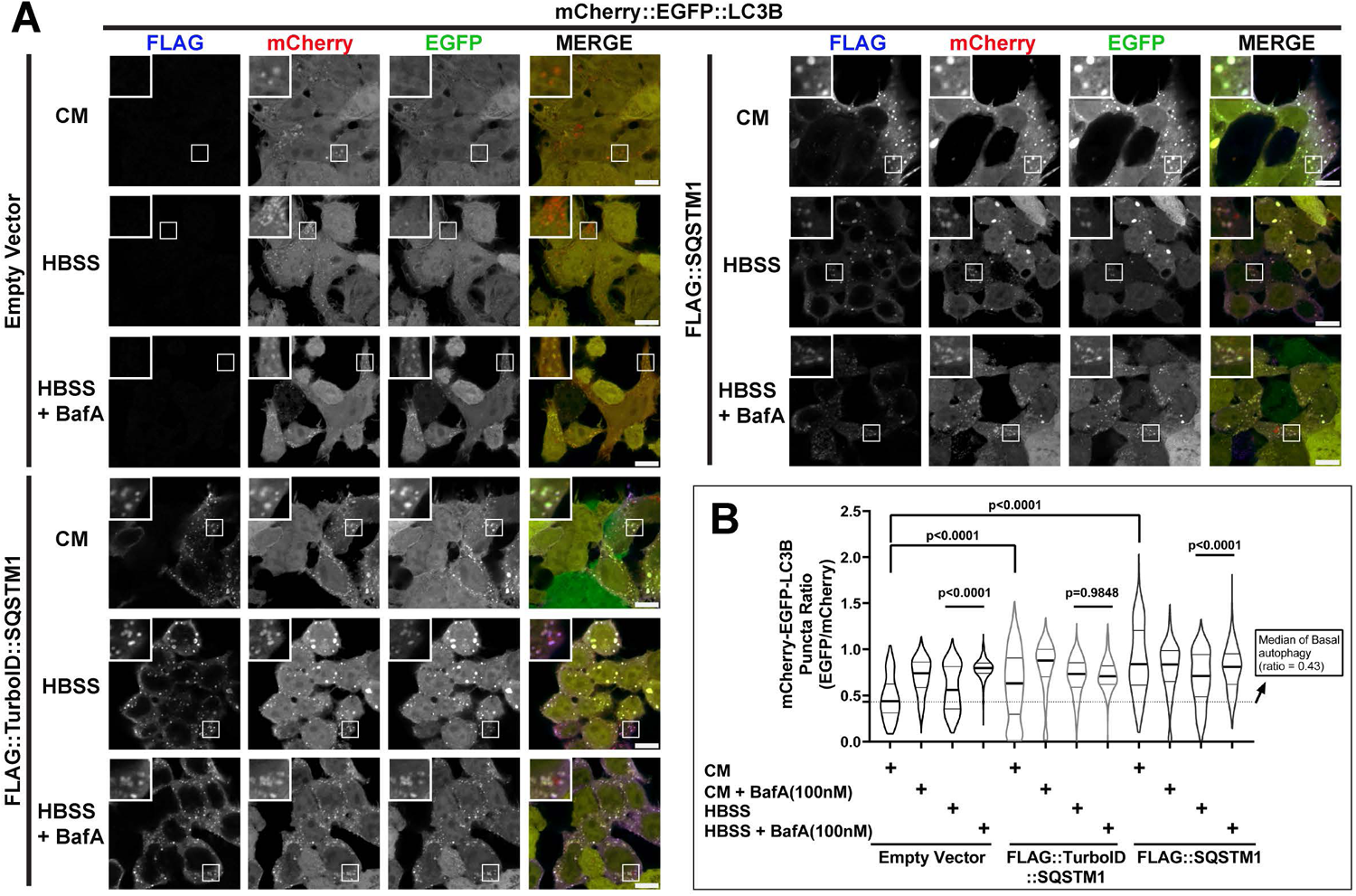
TurboID::SQSTM1 ovexpression inhibits autophagic flux. A) HEK-293T cells, expressing the autophagy flux sensor mCherry::EGFP::LC3B, were transduced with either Empty Vector, FLAG::TurboID::SQSTM1 or FLAG::SQSTM1 (MOI of 2). Fluorescence imaging of basal autophagy (CM), starved cells (HBSS), and autophagy inhibition (HBSS + BafA) were taken using confocal microscopy (scale bar = 10µm). B) Quantification of the puncta-ratio of EGFP/mCherry demonstrating that overexpression of both FLAG::TurboID::SQSTM1 and FLAG::SQSTM1 inhibits autophagy flux by sequestering LC3B into the SQSTM1 bodies. Statistical significance was determined using one-way ANOVA followed by a Tukey’s test. P-value < 0.05 was considered significant. Images contained (n=4) with at least 3 fields per condition, and the Data is listed in Supplementary Table 1, Sheet-9. CM, complete media; HBSS, Hanks’ balanced salt solution; BafA, Bafilomycin A1.

### Acute proteasomal inhibition modifies the SQSTM1 protein network

SQSTM1 facilitates degradation of ubiquitinylated proteins (42, 43). To assess how the SQSTM1 interactome changes with ubiquitin enrichment, we investigated the interaction network of FLAG::TurboID::SQSTM1 ± proteasomal inhibition using the small molecule inhibitor MG132 (44). FLAG::TurboID::SQSTM1-transduced HEK-293T cells were co-treated with biotin and either MG132 (50µM,) or vehicle (DMSO). After 2 hrs treatment, whole cell protein lysates were analyzed by Western Blotting. As expected, ubiquitinylated protein signals were elevated in the cells treated with MG132 (Figure 4A), consistent with inhibition of the ubiquitin proteasome degradation system (UPS).

**Figure 4.**
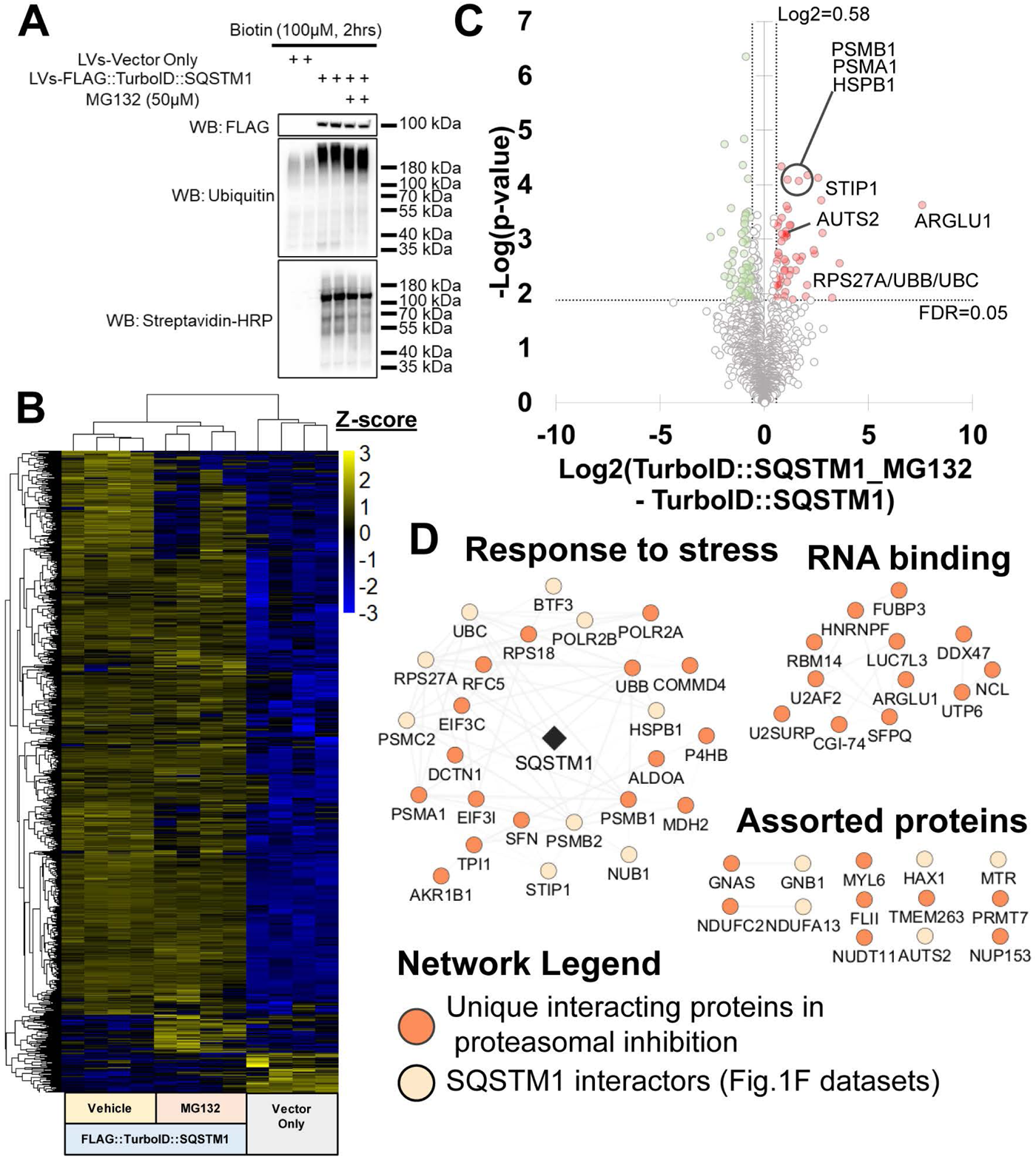
Acute proteasomal inhibition alters SQSTM1 protein network. A) MG132 inhibition of proteasome in HEK-293T cells, expressing TurboID::SQSTM1, induced higher levels of ubiquitinylated proteins by Western blot . B) Hierarchical clustering by Z-score intensities of differentially selected proteins in transduced HEK-293T cells. Proteins were selected using ANOVA and permutation-based FDR <0.05 (n=4). C) Quantitative comparison of -HEK-293T cells transduced with TurboID::SQSTM1 in presence or absence of MG132. Using Log2 FC(TurboID::SQSTM1_MG132 – TurboID::SQSTM1) of 0.58 and FDR < 0.05 to be considered significant (n=4). Data listed in Supplementary Table 1, Sheet-3. D) Protein network of significantly enriched proteins from Fig. 3C. Network was subclustered using functional annotation terms. Network legend displays unique SQSTM1 interacting proteins during proteasomal inhibition. FDR, False Discovery Rate; FC, Fold-Change.

Next we used streptavidin affinity-purification followed by LC-MS/MS to determine how the SQSTM1 protein interaction network (i.e., biotinylated proteins labeled by FLAG::TurboID::SQSTM1) changes in the context of proteasomal inhibition. Hierarchical clustering of the enriched proteins indicated that FLAG::TurboID::SQSTM1 treated with MG132 labeled different proteins compared to vehicle alone (Figure 4B). Using a criterion of Log2 FC (TurboID:SQSTM1_MG132 – TurboID::SQSTM1) > 0.58 and FDR < 0.05, 51 proteins including ubiquitin were significantly enriched from the MG132-treated cells (Figure 4C and Supporting Information-1, Sheet-3).

To assess the functional properties of this differential protein subnetwork, we cross-mapped annotations from the String-database. Subsequent visualization revealed 3 functional modules, including “Response to stress” and “RNA binding” (Figure 4D). These results demonstrated that TurboID::SQSTM1 is capable of capturing perturbations within the SQSTM1 interactome when proteostasis is altered.

### Tau aggregation alters SQSTM1 protein network in CRL-3275 cells

SQSTM1 protein is an autophagy receptor that facilitates degradation of misfolded proteins and protein aggregates. The aggregation-prone protein tau, encoded by the *MAPT* gene, has been found to interact with SQSTM1 in both human brain tissue with tauopathies and in murine transgenic tauopathies models (45–47). SQSTM1 overexpression has been found to ameliorate tau pathology in murine models (35). Given that FLAG::TurboID::SQSTM1 is capable of recognizing changes in the SQSTM1 interactome when proteostasis is dysregulated, we decided to determine how the presence of tau aggregates perturbs the SQSTM1 protein network. To achieve this, we used the well-established Tau RD P301S FRET Biosensor CRL-3275 cell line (CRL-3275 cells), which emits a FRET signal upon exposure to tau fibrils (Figure 5A) (48). The CRL-3275 cells were transduced with FLAG::TurboID::SQSTM1 and then treated with recombinant human P301S-MAPT protein fibrils (500ng/mL) or vehicle control for 24 or 48 hours; response of the CRL-3275 cells to tau aggregates was evident by imaging (Figure 5B). Transduced-CRL-3275 cells displayed both FLAG::TurboID::SQSTM1 expression and biotinylated protein signals associated with the FRET signal in the presence of tau fibrils, as determined by immunofluorescence and orthogonal projections of the images (Figures 5B and 5C).

**Figure 5.**
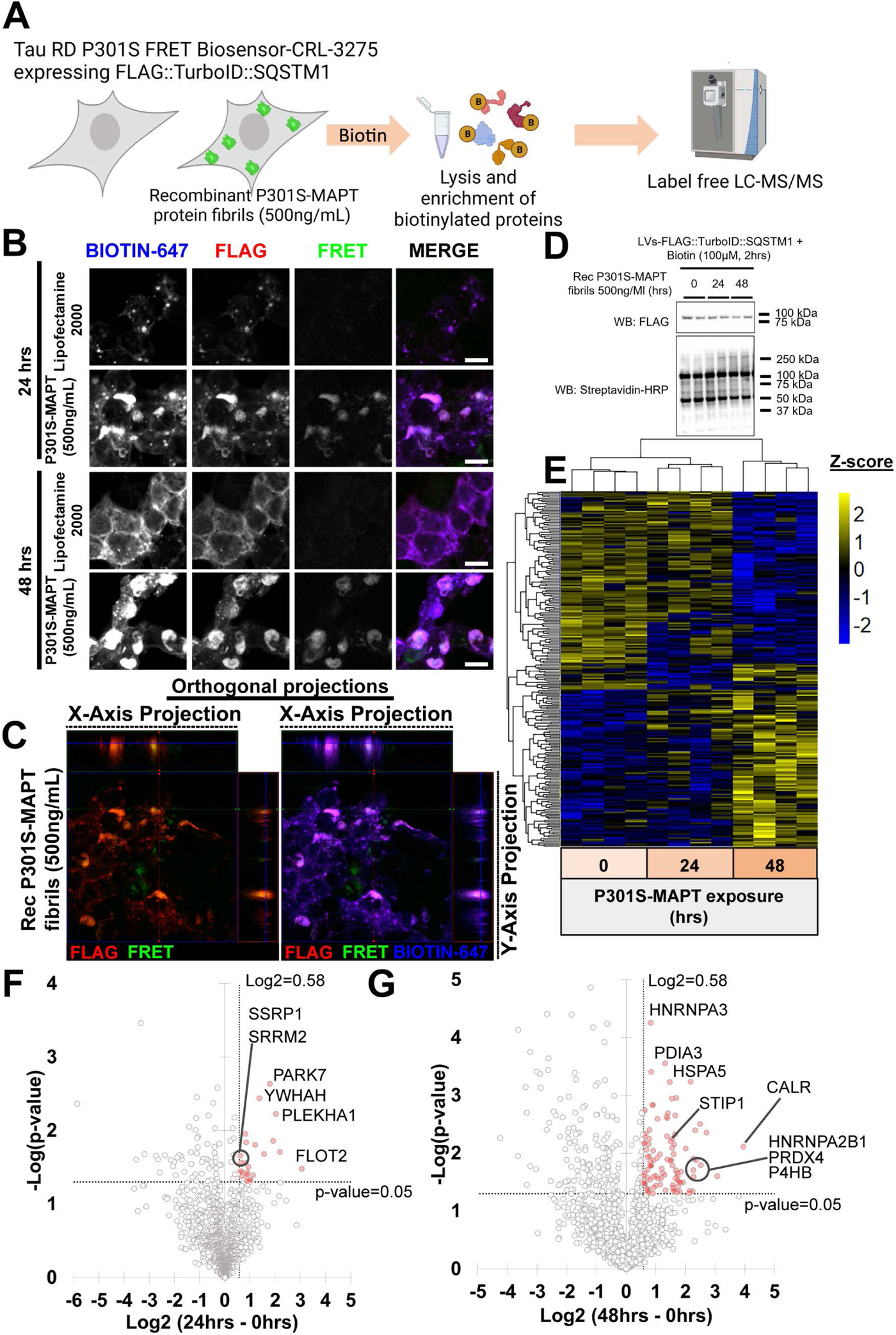
Tau aggregation alters SQSTM1 protein networks in CRL-3275 cells. A) Outline of the targeted proteomic approach of CRL-3275 cells, expressing TurboID::SQSTM1, in presence or absence of recombinant P301S-MAPT protein fibrils. B) Immunofluorescence of FLAG, biotinylated proteins and FRET signal from CRL-3275 cells transduced with TurboID::SQSTM1 treated with recombinant P301S-MAPT protein fibrils for 24 and 48 hours. C) Orthogonal projections of CRL-3275 cells, expressing TurboID::SQSTM1, treated with recombinant P301S-MAPT protein fibrils for 48 hours. Both FLAG and biotinylated protein signals co-localized with FRET signal. D) Western blot of TurboID::SQSTM1 transduced-CRL-3275 cells treated with recombinant fibrillar tau for 24 and 48 hours. E) Hierarchical clustering by Z-score intensities of differentially selected proteins in TurboID::SQSTM1 transduced-CRL-3275 cells and treated with recombinant P301S-MAPT protein fibrils for 24 and 48hrs. Proteins were selected using ANOVA and a p-value <0.05 (n=4). Data listed in Supplementary Table 1, Sheet–4. F and G) Quantitative comparison of CRL-3275 cells transduced with TurboID::SQSTM1 and treated with recombinant P301S-MAPT protein fibrils (24hrs vs 0hrs and 48hrs vs 0hrs). Using Log2 FC of 0.58 and p-value < 0.05 to be considered significant (n=4). Data listed in Supplementary Table 1, Sheets 5 and 6.

To explore the influence of tau aggregates on the molecular composition of the SQSTM1 protein networks, we transduced CRL-3275 cells with FLAG::TurboID::SQSTM1 and then treated them with recombinant fibrillar P301S-Tau protein (500ng/mL) for 24 and 48 hours prior to addition of biotin (100µM, 2hrs). Biotinylated proteins were detected by immunoblotting and then identified by LC-MS/MS (Figures 5E - and Supporting Information-1, Sheet-4). Hierarchical cluster analysis of labeled proteins indicated differential protein enrichment in the SQSTM1-protein network 48hrs post-treatment with fibrillar tau (Figure 5E). Volcano plots for the SQSTM1 network 24 or 48-hrs after treatment with fibrillar tau identified 24 and 98 proteins, respectively, exhibiting changes in SQSTM interactions (Log_2_ FC > 0.58, p-value < 0.05, Figures 5F and 5G - and Supporting Information-1, Sheets 5 and 6).

We proceeded to analyze the 98 proteins preferentially detected in the 48 vs 0 hr comparison. The SQSTM1 subnetwork induced by fibrillar tau was grouped based on functional annotations terms (Figure 6A – Supporting information-1, Sheet-8). We found that some of these enriched proteins are known MAPT interactors that were not detected in unstressed cells (Supplementary Table 1, Sheet-8). Notably, many of these MAPT interacting proteins are not reported as SQSTM1 interactors in existing datasets (BioGRID-SQSTM1), several of these proteins (*e.g.* P4HB, GPI, and ANXA5) are highlighted in Figure 6A (orange diamonds).

**Figure 6.**
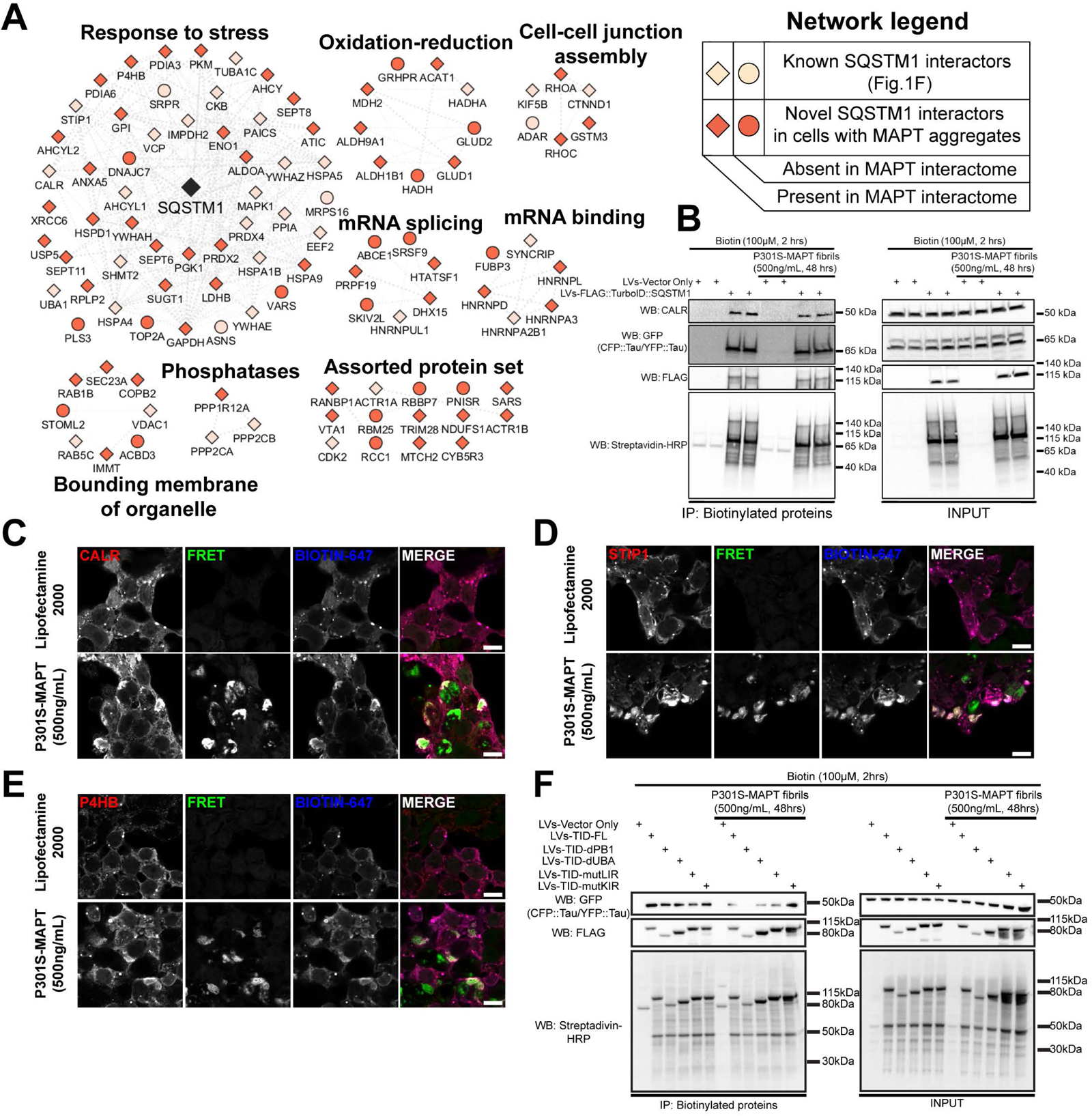
Tau aggregation sequesters SQSTM1 interacting proteins in CRL-3275 cells. A) Protein network of significantly enriched proteins from Fig. 5G. Note that SQSTM1 was not found significant in the dataset from Fig.5G. However, SQSTM1 was added in the network to display the interaction of TurboID::SQSTM1 with the protein network. Network was subclustered using functional annotation terms. Network legend displays unique SQSTM1 interacting proteins in presence of MAPT aggregates. Datasets used in the network legend: BioGRID-human-SQSTM1, BioGRID-human-MAPT, (47, 77, 78). B) Pull-down of biotinylated proteins of CRL-3275 expressing lentiviral constructs treated with recombinant MAPT fibrils for 48hrs. Western blot indicates interaction of TurboID::SQSTM1 with CALR and Tau domains (CFP::K18-tau and YFP::K18-tau). C, D and E) Immunofluorescence of CALR, STIP1 and P4HB in TurboID::SQSTM1-transduced CRL-3275 cells in presence of recombinant fibrillar tau for 48hrs. F) Pull-down of biotinylated proteins of CRL-3275 expressing different TurboID::SQSTM1 iterations (Fig. 2E) treated with recombinant MAPT fibrils for 48hrs. Western blot indicates K18-tau aggregates requires PB1 domain from SQSTM1 to interact with TurboID::SQSTM1 in presence of MAPT fibrils for 48hrs.

### Tau aggregation sequesters SQSTM1-interacting proteins in CRL-3275 cells

Calreticulin (CALR) is a chaperone protein that resides in the endoplasmic reticulum (ER). This protein has been reported to interact with SQSTM1 in a process called ER-phagy by two mechanisms: a) an indirect interaction via the ER E3-ligase TRIM13 and b) direct interaction as an N-recognin (49, 50). Strikingly, CALR was the most abundant biotinylated protein, but showed similar levels of biotinylation ± fibrillar tau (Figure 5G). We independently tested interaction of SQSTM1 with CALR by pulling-down biotinylated proteins from CRL-3275 and immunoblotting (Figure 6B). Both CALR and tau domains (CFP::K18-tau and YFP::K18-tau) were successfully pulled-down, but the interactions were not affected by the presence of fibrillar tau.

Although fibrillar tau did not change the amount of CALR interacting with SQSTM1, we hypothesized that fibrillar tau might change the subcellular localization of the SQSTM1-CALR interaction. The intracellular localization of CALR was imaged by immunofluorescence in FLAG::TurboID::SQSTM1-expressing CRL-3275 cells in presence or absence of recombinant tau fibrils (Figure 6C). Strikingly, CALR showed areas of consolidation in cells exposed to tau fibrils; these areas of consolidation co-localized with both FRET signal (tau aggregation) and biotinylated proteins (FLAG::TurboID::SQSTM1) indicating the association of CALR, SQSTM1 and tau aggregates (Figure 6C). Thus, aggregation of tau leads to sequestration of SQSTM1, which also leads to sequestration of CALR.

To broaden our assessment, we examined whether tau aggregation sequesters other stress-induced SQSTM1 protein subnetwork members. We proceeded to evaluate the cellular localization of two other cellular proteins, STIP1 and P4HB, each of which exhibited significantly higher association with SQSTM1 in presence of fibrillar tau at 48 hrs vs 0 hrs (Figure 5G). STIP1 was previously proposed to interact with SQSTM1 (29) and was also identified in our dysregulated-proteostasis SQSTM1 protein network (Figure 4C). P4HB was previously described to interact with MAPT (51). Both proteins exhibited pronounced changes in intracellular localization in response to the accumulation of aggregated tau, as was observed with CALR (Figures 6C-E). Importantly, the subcellular patterns of STIP1 and P4HB followed SQSTM1 sequestration in response to the accumulation of tau fibrils and co-localized with the resulting FRET signal (tau aggregation, Figures 6D and 6E).

To identify whether a particular functional domain of SQSTM1 interacts with aggregated K18-Tau peptide, we transduced CRL-3275 cells with the TurboID::SQSTM1 dysfunctional-domains constructs, and then treated the cells with P301S-MAPT fibrils (500ng/mL for 48hrs). Next, biotinylated proteins were pull-down and interaction of SQSTM1 with the K18-tau peptide was monitored by Western blot (CFP::K18-tau and YFP::K18-tau). The results demonstrate that deletion of PB1 domain abrogated the interaction aggregated K18-tau peptide with SQSTM1 (Figure 6F).

Collectively, these experiments demonstrate that SQSTM1 proximity labeling identifies changes in the SQSTM1 protein interaction network in responses to tau aggregation and enables discovery of novel proteins that associate dynamically with the SQSTM1 response.

## DISCUSSION

In the current study, we generated chimeric TurboID::SQSTM1 constructs consisting of full length protein, domain deletion constructs or domain mutation constructs (Figure 2E). These chimeric proteins enable proximity labeling of the SQSTM1 protein interaction network by combining TurboID with full-length human SQSTM1 protein. Using targeted proteomics, the TurboID::SQSTM1 chimeric protein uncovered components of the dynamic SQSTM1 protein network in homeostatic and stress conditions, resulting in the identification of both known and novel SQSTM1 interacting proteins (Figures 1 and 2). The novel SQSTM1 interacting partners include proteins associated with dysregulated cellular-proteostasis or the presence of tau protein aggregates (Figures 4 and 5).

SQSTM1 plays a central role in cellular catabolism by engaging with diverse cellular proteins that are targeted for degradation and shuttling them to the autophagosome. This broad ability to identify substrate binding and intracellular aggregates positions the TurboID::SQSTM1 chimera as an informative tool for interrogating macromolecular changes in the composition and evolution of protein aggregates as they accumulate in disease model systems (21, 22, 52). The TurboID::SQSTM1 chimera also serves as a useful resource to broadly explore SQSTM1 functions in an unbiased manner (21, 22, 52). The SQSTM1 protein links ubiquitinylated proteins to autophagosome for degradation (26, 40), forms biomolecular condensates (42, 53, 54), and interacts with small non-coding RNAs to regulate selective autophagy (55). The TurboID::SQSTM1 chimera complements other proximity labeling resources that by enabling comprehensive analysis of SQSTM1 interacting proteins *in vitro* and *in vivo* (6, 29). Use of the TurboID::SQSTM1 chimera will particularly facilitate studies investigating how the SQSTM1-dependent network changes within disease models.

The ability to identify known SQSTM1 interaction partners with TurboID::SQSTM1 proximity profile labeling suggests that addition of TurboID to SQSTM1 construct preserves function of the different SQSTM1 protein domains (PB1, UBA, LIR and KIR). Importantly, the perturbations of these essential SQSTM1 functional domains impaired the interaction with known SQSTM1-interacting partners, as observed in Figures 1 and 2. This important result suggests that the interactions of the TurboID::SQSTM1 chimeric protein will reflect the normal biology of SQSTM1 when applied to *in vivo* and *in vitro* systems. TurboID::SQSTM1 also identified novel SQSTM1 interacting-proteins, including proteins that are associated with translation (Figure 1F). Some of these translation-associated proteins are RNA-binding proteins that contain intrinsically disordered regions (IDRs). Our dataset identified AGO2, which contains an IDR and is a member of the RNA-induced silencing complex (Figures 1 and 2). The association of RNA binding proteins such as AGO2 with SQSTM1 provides a putative mechanism for the presence of RNA in SQSTM1 bodies and might contribute to the tendency of SQSTM1-bodies to form biomolecular condensates (42, 53). SYNRG is another novel protein described in our dataset. SYNRG functions as an adaptor protein between AP-1 to clathrin-associated proteins that potentially mediates the association of SQSTM1 with β-arrestin (9, 10, 56) and cannabinoid receptor-2 (28) Thus, novel proteins labeled by TurboID::SQSTM1 provide important new insights about SQSTM1 biology.

The chimeric protein TurboID::SQSTM1 successfully identified perturbations in the SQSTM1 protein network composition during proteasomal inhibition (Figure 4). UPS inhibition induces the translational stress response, which is known to alter the function and distribution of RNA binding proteins (57). Applying the TurboID::SQSTM1 protein in the context of UPS inhibition identified multiple splicing-associated RNA binding proteins, as shown in the sub-set of proteins labelled as ‘RNA binding’ including DDX47, FUBP3, SFPQ and HNRPF (Figures 4C and 4D). These findings are consistent with previous results indicating that UPS inhibition impacts on alternative splicing and affects the generation of protein variants (58). Our datasets identified an interaction between ARGLU1 and SQSTM1 that occurs selectively in response to proteasomal inhibition (Figures 4C and 4D). Interestingly, ARGLU1 was previously found to be an essential transcriptional co-activator and splicing regulator of glucocorticoid receptor (59). ARGLU1 and 5 other genes were recently found to be applicable as biomarkers for amyotrophic lateral sclerosis, using weighted gene co-expression network analysis (60). These results show the utility of TurboID::SQSTM1 protein as a tool for detecting perturbations in SQSTM1 network, using acute UPS inhibition as a model. This tool also demonstrates its capability to interrogate cellular stress mechanisms or disease processes.

SQSTM1 protein facilitates the degradation of protein aggregates by linking ubiquitinylated proteins to autophagy (1, 35, 61). Our findings demonstrate how SQSTM1 protein networks are altered by the presence of a protein aggregates, such as recombinant fibrillar tau (Figures 5 and 6). The protein networks in both Figure 4 and Figure 5 highlight the pathway ‘Response to stress’, which includes many chaperones commonly associated with misfolded proteins, including both cytoplasmic chaperones (STIP1, HSPB1, HSPA4, HSPA1B, etc.) and ER resident chaperones (CALR, P4HB, HSPA5 and others). The co-chaperone STIP1 is a stress-induced gene that has an essential role presenting protein substrates to HSP90. A proteomic dataset recently reported that the STIP1-SQSTM1 interaction is elevated during autophagy inhibition (29). STIP1 was also recently proposed to be a potential druggable target for neurodegeneration (62). These results show how the TurboID::SQSTM1 protein can be used to identify SQSTM1 interactors that function in protein degradation and the ‘Response to stress’ pathways (29).

The interaction of tau protein and SQSTM1 has been widely described (14, 63, 64). This interaction is proposed to occur through interaction of ubiquitinylated tau with UBA domain from SQSTM1 (46), and direct interaction of ZZ domain of SQSTM1 with N region of tau protein via tau proteolysis by N-terminal arginylation (49, 65). In this report we investigated the perturbation of SQSTM1 network by tau aggregation using CRL-3275 cells and identified that PB1 domain of SQSTM1 is necessary for interaction with aggregated K18-tau peptide (48, 66). The resulting networks identify multiple known MAPT protein interactors that are recognized by TurboID::SQSTM1 as part of the SQSTM1 network, including CALR, HNRNPA2B1, HSPA5, etc. (47, 67). The TurboID::SQSTM1 dataset also describes novel SQSTM1 interactors that appear only in the presence of tau aggregates (*i.e.* P4HB, SYNCRIP, DNAJC7; Figure 5A). Prior studies described multiple SQSTM1 interactors interacting with tau aggregates including LC3B, UBB, UBA52, and CALR (45–47). The incomplete overlap between existing datasets and our results might reflect a difference between the relatively simple biology of a HEK-293T cell versus the brain. Focusing on SQSTM1 networks rather than tau networks might also direct attention towards proteins that are in the aggregate but associate more closely with SQSTM1 than with tau protein. Thus, this work highlights potentially preferential interactions among interactors of SQSTM1 and tau.

In summary, we provide this novel and validated resource to explore SQSTM1 protein network. TurboID has the capability to explore protein networks *in vitro* and *in vivo,* TurboID::SQSTM1 can then be used in a context-dependent manner to explore SQSTM1 networks (studying tauopathies murine models, cancer research, other neurodegeneration models, cellular-specific SQSTM1 network behavior, etc.). TurboID::SQSTM1 can also identify novel proteins that can be exploited as novel therapeutic targets (*i.e.* AUTOTAC and PROTAC technologies).

## LIMITATIONS

TurboID::SQSTM1 offers a broad view of the SQSTM1 protein network. This chimeric protein is an overexpressed system that might affect the protein-protein interactions stoichiometry. For that reason, it is important to further validate the novel proteins with alternatives approaches (*e.g.* immunoprecipitation, proximity ligation assay, surface plasmon resonance, microscale thermophoresis, etc.). The present study presents novel SQSTM1 interactors that can be useful for autophagy research (and protein degradation in neurodegenerative disorders). However, the biological mechanisms of these novel interactors remain to be explored to understand their roles in proteostasis.

## EXPERIMENTAL PROCEDURES

### DNA constructs

We generated the DNA construct FLAG::TurboID::SQSTM1 by amplifying human SQSTM1 from Dr. Qing Zhong’s plasmid (Addgene#280227(68)) and TurboID from Dr. Alice Ting’s plasmid (30) via PCR using Q5 high-fidelity (M0492, NEB). Next, we cloned both amplicons into the pHR-SFFV lentiviral vector (Addgene#79121, (69)) using In-Fusion cloning kit (Takara). Truncated versions of FLAG::TurboID::SQSTM1 were generated by PCR using primers corresponding to the truncated SQSTM1 gene and amplification using Q5 high-fidelity (M0492, NEB). Loss-of-function mutations were generated using site directed mutagenesis using Q5 Site-directed mutagenesis kit (E0552S, NEB) with the HA::SQSTM1/p62 plasmid as template (Addgene#280227). We verified the sequences of all lentiviral constructs using Sanger sequencing. These constructs can be found on Addgene.

### Mammalian cell culture and Lentiviral packaging

We maintained HEK-293T (CRL-3216, ATCC) and Tau-RD-P301S-K18 FRET Biosensor CRL-3275, ATCC, (48) cell lines in DMEM supplemented with 10% FBS and 1% penicillin/streptomycin. The cells were grown in plates coated with poly-d-lysine (PDL) and in a humidified environment with 5% CO2. To package lentivirus, we grew HEK-293T cells in PDL-coated 10-cm petri dishes and transfected them with transfer plasmids, psPAX2 (Addgene#12260), and VSV-G (Addgene#8454) at a ratio of 1:1:3, respectively, using Fugene-HD reagent (Promega E2311). Three days later, we harvested and concentrated the lentiviral particles using the LentiX concentrator (Takara #631232) following the manufacturer’s protocol. The lentiviral titer was then determined using the Lenti-X p24 rapid titer kit (Takara Cat #632200). The Tau-RD-P301S-K18 FRET Biosensor CRL-3275 were exposed to active recombinant P301S-MAPT fibrils (ab246003) as described by the Diamond laboratory (48)

### Transduction and Transfection

To perform the experiments, HEK-293T and CRL-3275 cells were grown in PDL-coated 6-well plates and transduced with lentiviral particles using a multiplicity of infection value of 2. Subsequently, these cells were passaged into either PDL-coated cover slips, 6-well plates, or 10-cm petri dishes. For HA-LC3B-Myc (Addgene #137757), hrGFP-KEAP1 (Addgene #28025) and HA-SQSTM1 (Addgene #280227) transfections, HEK-293T cells were transfected using Lipofectamine 3000 (Thermo Fisher), following the manufacturer’s protocol.

### Biotin labeling, Immunocytochemistry, Immunoblot and Biotinylated proteins-streptavidin pull-down

Transduced Hek-293T cells were treated with biotin (100µM, 2 hours). Next, these cells were placed on ice for 10 minutes. Subsequently, cells were fixed with 4%PFA in PBS for 15 minutes. Fixed cells were permeabilized using ice-cold methanol for 5 minutes, as previously described (22). Cells were then blocked with 5% donkey serum, 5% bovine-serum albumin and 22.52 mg/mL of glycine in PBS-T. Antibodies are described in Table 1.

**Table 1.** Antibodies used in this study.

| Application | Antibody | Host | Manufacturer | Cat number | Dilution |
| --- | --- | --- | --- | --- | --- |
| ICC | FLAG | Mouse | Sigma | F1804 | 1:200 |
| WB | Streptavidin-HRP | N/A | Thermo Fisher | 21126 | 1:5000 |
| WB | Ubiquitin | Mouse | Cell Signaling | 3936 | 1:2000 |
| WB | FLAG | Mouse | Sigma | F1804 | 1:2000 |
| WB | GFP | Rabbit | Thermo Fisher | A-11122 | 1:5000 |
| IP/WB | KEAP1 | Rabbit | Cell Signaling | 8047S | 1:1000 |
| IP/WB | SQSTM1 | Rabbit | ABCAM | ab109012 | 1:10000 |
| ICC | HA | Mouse | BIOLEGEND | 901502 | 1:250 |
| WB/IP | HA | Mouse | BIOLEGEND | 901502 | 1:2000 |
| ICC | LC3B | Rabbit | ABCAM | AB192890 | 1:100 |
| WB | CALR | Rabbit | ABCAM | ab92516 | 1:5000 |
| wb | CALR | Rabbit | ABCAM | ab92516 | 1:5000 |
| ICC | STIP1 | Rabbit | ABCAM | ab126724 | 1:10000 |
| WB/IP | P4HB | Rabbit | ABCAM | ab137110 | 1:2000 |
| WB/IP | AGO2 | Rabbit | ABCAM | ab156870 | 1:2000 |
| WB/IP | SYNRG | Rabbit | BETHYL<br>BIOSCIENCES | A304-567A | 1:2000 |
| WB | Anti Mouse IgG (H+L)<br>Secondary Antibody,<br>HRP | Goat | Thermo Fisher | 62-6520 | 1:5000 |
| WB | Anti Rabbit IgG (H+L)<br>Secondary Antibody,<br>HRP | Goat | Thermo Fisher | 62-6120 | 1:5000 |
| ICC | Streptavidin-DyLight<br>488 | N/A | Vector Labs | SA-5488-1 | 1:2000 |
| ICC | Streptavidin-DyLight<br>649 | N/A | Vector Labs | SA-5649-1 | 1:2000 |
| ICC | Alexa Fluor conjugated<br>secondary antibodies | Donkey | Jackson<br>Immunoresearch |  | 1:1000 |

Following biotin labeling, cells were rinsed with ice-cold PBS twice and lysed in RIPA buffer (0.5% sodium deoxycholate, 1% TritonX100, 0.5% SDS, 1mM EDTA, 150mM NaCl, 50mM Tris-HCl pH 7.5). Protein concentration was determined using BCA assay and 40µg of total proteins were resolved using Western blot. For streptavidin pull-downs, 3 mg of total proteins were pulled-down using Pierce Streptavidin magnetic beads (Cat#88816).

### Confocal microscopy and Autophagy flux assessment

Immunofluorescence images were taken using a Zeiss LSM700 laser-scanning confocal microscope (63X oil objective). Pinhole size was kept at 1 Airy Unit and images were obtained using both Zen Black and Zen Blue Softwares from Zeiss, as previously reported (67, 70).

Autophagy flux analysis was performed using FIJI/ImageJ (NIH). Briefly, images were taken using mCherry, EGFP and near infrared fluorescence channels (Alexa Fluor 647 for FLAG-tagged proteins). Then, raw pictures were uploaded into FIJI and mCherry puncta from mCherry::EGFP::LC3B autophagy sensor was to create puncta mask. Following, the mCherry puncta mask was used to determine mean gray fluorescence intensity of mCherry and EGFP in each puncta per condition. Moreover, the puncta-ratio was used to determine the autophagy flux. The EGFP/mCherry puncta-ratio values from each mCherry-masked puncta are found in the Supporting Information-1, Sheet-9.

### Immunoprecipitation

HEK-293T cells transiently expressing HA::SQSTM1 were lysed in IP-buffer (0.1% NP-40, 1mM EDTA, 120 mM NaCl, 50mM tris-HCl pH 7.5) on ice. Protein concentration was determined by BCA assay and 2 mg of total protein was taken. Following, 2µg of anti-HA antibody was added to each sample and incubated for 2 hours at 4C. Subsequently, 50µL of Dynabeads Protein G (ThermoFisher Cat#10004D) were added to each sample and the samples were incubated overnight at 4C. Next day, the samples were washed with IP-buffer 3 times (5 minutes each) and samples were eluted in WB loading buffer.

### Sample preparation for LC-MS/MS

After cell transduction, cells expressing either FLAG::TurboID or FLAG::TurboID::SQSTM1 were plated in PDL-coated 10-cm dishes. Upon reaching confluency, a subset of the cells was treated with 100 µM biotin 2hrs to induce protein biotinylation. The cells were subsequently lysed in Lysis buffer containing 50mM Tris-HCl (pH 7.5), 150mM NaCl, 1mM EDTA, 0.5% SDS, 1% Triton-X100 and 0.5% sodium deoxycholate. The lysis buffer was supplemented with protease inhibitors (Halt protease inhibitor cocktail, Thermo Fisher # 78429). Protein levels were quantified using a BCA assay kit from Thermo Fisher. Equal protein amounts (3mg) were immunoprecipitated overnight at 4C using Streptavidin Magnetic Beads (Pierce #88816). Following the streptavidin pull down, the beads were washed twice with RIPA buffer, once with KCl 1M, once with Na2CO3 0.1M, once with Urea 2M (in 10mM Tris-HCl, pH 8) and twice with ammonium bicarbonate 50mM as previously described (22)

To perform on-bead digestion, the beads were suspended in 200 µL of 50 mM ammonium bicarbonate and incubated for 30 minutes at room temperature with 40 mM chloroacetamide and 10 mM TCEP to alkylate and reduce the proteins, respectively. The reaction was then quenched using 20 mM DTT. The beads were subsequently digested overnight at 37C with 1 µg of MS-grade trypsin from Thermo Fisher Scientific (Cat#90058). The next day, the digested peptides were collected into new tubes, and the trypsin-digestion reaction was quenched with formic acid (final concentration of 1%). These peptides were then desalted using Pierce C18 columns (Thermo Fisher Scientific, Cat#89870), following the manufacturer’s protocol, and finally stored at -80C for further proteomics analysis.

### HPLC-ESI MS/MS and Data Analysis

The LC-MS/MS analysis was performed as previously described (67, 71). In brief, the desalted peptides were reconstituted in 1% formic acid and fractionated using C18 PepMap pre-column (3µm, 100Å, 75mm x 2cm) hyphenated to a RSLC C18 analytical column (2mm, 100 Å, 75 µm x 50cm) High performance nanoflow liquid chromatography-Orbitrap tandem mass spectrometry (LC-MS/MS) were performed using the Easy nLC 1200 system coupled to Q-Exactive HF-X MS (Thermo Scientific).

LC-MS/MS raw data were processed in MaxQuant v1.3.7.0 (72), using the Human proteome (2018_04 Uniprot release of UP000005640_9606) with match between-runs activated. Then, the MaxQuant output file was analyzed in Perseus v2.0.3.0 (73) using a modified version of the settings described by the Long Lab (24) Briefly, LFQ intensity protein values from all conditions were Log_2_ transformed, proteins with at least three quantified values per condition were retained, and missing values were replaced using imputation processing from a normal distribution with a width of 0.3 and a downshift value of 1.8 (these are the default values). Statistical comparisons were performed in Perseus applying two-tailed *t-*test with either a permutation-based FDR of 0.05 or *p-*value less than 0.05. ANOVA was used for a two-dimensional hierarchical clustering considering a permutation-based FDR of 0.05.

Enriched proteins from the comparisons were cross-referenced using the interactome database BioGRID (accessed on 11.21.2021). Additionally, these enriched proteins were used to build protein networks in Cytoscape Version 3.9.1 and this network was sub-clustered with the ClusterMaker2 Version 2.3.2 plugin. Functional annotation terms were determined using ShinyGO Version 0.76.3 (74).

## DATA AVAILABILITY

All obtained raw data have been deposited to the ProteomeXchange Consortium via the PRIDE partner repository with the dataset identifier PXD047725 (75, 76). All constructs designed in this study will be deposited in Addgene.

PRIDE access:

**Project Name:** Proximity labeling reveals dynamic changes in the SQSTM1 protein network

Project accession: PXD047725

**Project DOI:** Not applicable Reviewer account details:

Password: eda2jKGx

